# An Integrated Modelling Framework for the Forest-based Bioeconomy

**DOI:** 10.1101/011932

**Authors:** Sarah Mubareka, Ragnar Jonsson, Francesca Rinaldi, Giulia Fiorese, Jesús San-Miguel-Ayanz, Ola Sallnäs, Claudia Baranzelli, Roberto Pilli, Carlo Lavalle, Alban Kitous

## Abstract

This paper describes the conceptual design of a regional modelling framework to assess scenarios for the forest-based bioeconomy. The framework consists of a core set of tools: a partial equilibrium model for the forest sector, a forestry dynamics model for forest growth and harvest and a wood resources balance sheet. The framework can be expanded to include an energy model, a land use model and a forest owner decision model. This partially integrated, multi-disciplinary modelling framework is described, with particular emphasis on the structure of the variables to be exchanged between the framework tools. The data exchange is subject to a series of integrity checks to ensure that the model is computing the correct information in the correct format and order of elements.

## Introduction

### Bioeconomy

The “bioeconomy” refers to the sustainable production and conversion of biomass into a range of industrial and non-industrial products and energy. It offers opportunities and solutions to a growing number of environmental and economic challenges, including the management of natural resources, the mitigation of climate change and the provision of material goods, energy and food. The forest-based sector is instrumental for the successful implementation of the EU Bioeconomy Strategy [1]. It can contribute to mitigate climate change, both through carbon sequestration in forests, as well as in harvested wood products; and through the substitution of carbon-intensive materials such as fossil fuels or cement. The forest sector interacts with both the energy and industrial products sectors among others, therefore requiring a systems approach to this analysis [2]. An integrated and coordinated approach is thus required to assess both the full potential and the impacts of the bioeconomy. The various interdependent sectors of the bioeconomy must be properly linked through coordinated sectoral components in order to form a multi-disciplinary modelling framework.

The purpose of the proposed integrated modelling framework is to generate a periodical analysis on the availability of biomass from forests at regional level in the EU; and the present and future level of sustainability for the provision of this raw material under different scenarios for different regions. Scenarios may be based on different demand settings; or may be based on the provision itself, such as policies affecting wood mobilisation. The system is designed to answer questions that arise concerning the impacts and potential of the bioeconomy on the EU’s forest sector at a regional level. In order to do this, we need tools to measure and forecast the EU wood production potential; wood product demands inside and outside the EU; availability of products outside of the EU; energy requirements from biomass; and effects of current and future policy, including those affecting the environment.

### Integrated modelling approach

Integrated modelling is “..the process of combining several sub-models that represent different interacting natural and social systems..” [3]. It is a useful approach used in multi-disciplinary quantitative analysis, to bring pieces of sector-specific models together to form a more comprehensive picture of system dynamics. This provides a platform for combining knowledge from diverse scientific disciplines. In this way, the effect of a sector-specific policy on a more comprehensive chain of events can be assessed. In an integrated modelling approach, models from different scientific disciplines are independently developed, which means that experts are responsible for their own specific modules.

Integrated assessment models are widely used as a multidisciplinary tool for policy evaluation. Different approaches can be used to integrate disciplinary models into one modelling framework. The integration or linking of disciplinary models, however, presents several challenges including maintaining the consistency between models that operate at different spatial and temporal scales [4]. Different approaches to address these challenges are reflected in the resulting degree of integration of the sectoral models within one integrated model [5, 6, 7]; the sectoral models can remain stand-alone e.g. Böttcher et al. [8] and van Ittersum et al. [9]] or be integrated into a single model (e.g. IMAGE [10]).

In this system, the choice is to maintain the integrity of the individual models, while facilitating the interaction between the models through bridges, or data transformation modules (D-TM). This approach has several advantages. First, the system remains modular, thus only the components that are necessary are active; second, individual models remain independent and within the control of domain experts, thus allowing for developers to improve upon their own models, whether for the benefit of the system or for other applications. A third benefit is the uniqueness of this type of system for Europe. Its individual components have been (or are currently being) developed within one research organization, thus facilitating repeatability and exchange between modellers.

### Semantic, conceptual and technical integration of models

Interoperability of sector-specific models means that these may be running on entirely different computer platforms. There must therefore be a shared understanding between disciplines to ensure the consistency and transparency in definitions and terms. Ontologies, concept maps, variable mapping are all used for this purpose [11, 12]. This implies that “real-world” issues to be represented and eventually solved through modelling, are formulated in the universal language of mathematics. Often however, a fundamental concept, such as the fact that an *array* of data should be processed, gets lost in the mathematical description of the individual model parameters, making dialogue between modellers less comprehensible. For example, the variables that interact between models may actually need to interact with a frequency of five-year time steps, updating the situation at every time step. This piece of information may get lost when describing, in mathematical form, the interaction between variables. Other points for misinterpretation, when dealing purely with mathematical formulation, are the scale of the analysis, common driving forces behind the models, spatial interactions, temporal dynamics, and even the degree to which the models should be integrated.

Integrity checks are responsible to stop the model if key variables are not in the expected format. This is even more important when several models are integrated. These types of integrity checks are only possible if semantic constraints are made because they offer an infrastructure of support in the design phase of model integration. In this way, the semantic properties of input and output are clear within a multi-disciplinary context regardless of discipline-specific annotations. This is relevant in the conceptual phase of the integration of models because it refers to the alignment of input/output between models. This paper addresses these issues within the multi-disciplinary modelling framework for modelling the forest sector within the context of the EU bioeconomy. In this paper we condense complex features into simple variables in order to facilitate the communication between the different disciplines. Each variable is derived from a series of calculations within the discipline’s own model.

## Methods

### General description of models

Table 1 gives an overview of the tools (models and datasets) that could be used for the assessment of the forest-sector related aspects of the bioeconomy. We have separated the tools into two groups: a core toolset and an extended toolset. The core set of tools is key to understanding the interaction between detailed forest resource availability and the global demands for forest products for both material and energy uses. These tools depend heavily on one another and are fully integrated and include a global dynamic partial equilibrium model for the forestry sector, the Global Forestry Trade Model (GFTM); a forestry dynamics model, a niche filled either by the European Forestry Dynamics Model (EFDM) or the Carbon Budget Model (CBM); and the Wood Resources Balance sheet (WRB), an accounting tool that provides a summary of the requirements and availability for forest products.

**Table 1:**
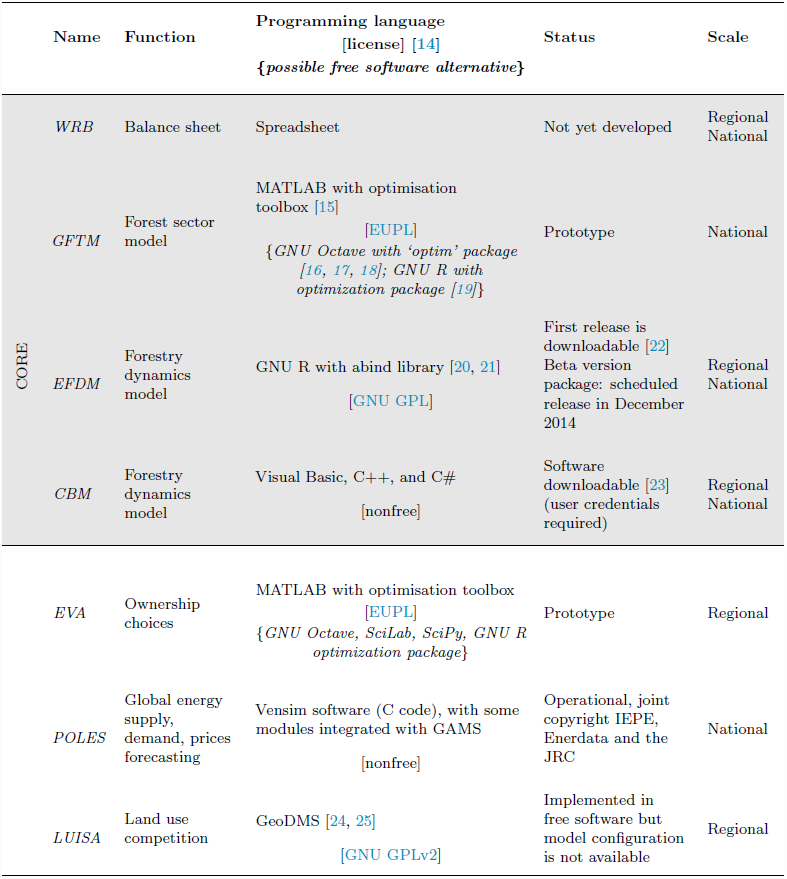
Summary of tools in assessing the forest-based sector of the bioeconomy. The core required tools are shaded in grey, while others are extensions of the framework.

In addition to the core set of tools, other tools can be used to improve the modelling platform. Models include the Expected Value Asymmetries (EVA) model for ownership behaviour; the energy sector model POLES (Prospective Outlook on Long-term Energy Systems^1^); and the Land-Use-based Integrated Sustainability Assessment Modelling Platform (LUISA [13]).

Throughout the development of the different models, the emphasis has been on the expert knowledge of the persons developing the model. Any effort to harmonise the programming language of the models is therefore avoided because control of the model should be maintained by the domain expert in the language they deem most suitable. The pieces of software that link the models together, mainly for data transformation and the integrity check, is written in free software (Python [26]).

As shown in Figure 1, the models interact iteratively. Loose, initial demand and supply constraints are generated to launch the computation process. These initial demand and supply values are then tightened in a second iteration.

**Figure 1.**
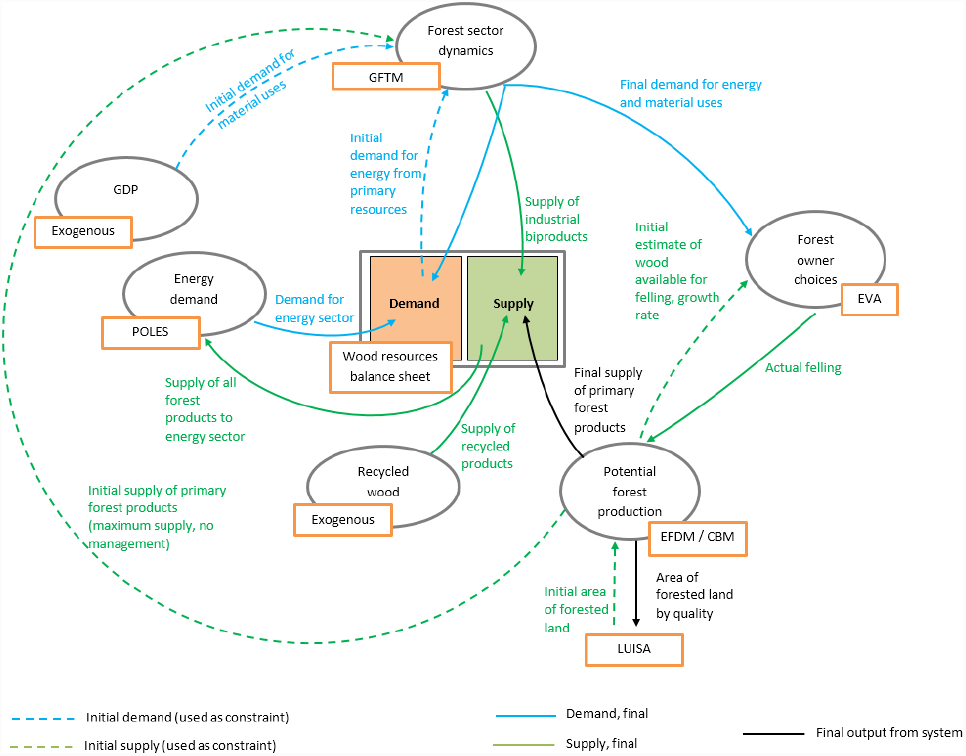
Overview of interaction between models for the forest-based sector in the bioeconomy.

### The core modelling system

Central to the forest-based bioeconomy modelling framework is the forest sector model, GFTM. GFTM estimates wood demand and supply given global drivers and the baseline internal availability of wood for a selected time span. GFTM is an equilibrium model for the forest sector that includes international trade and ensures balance between sources and uses of woody biomass for a given geographical unit. GFTM requires information about the total maximum availability of wood from EFDM.

EFDM belongs to the forest model family of matrix models, born in the 1940’s to model future plant and animal population structures. Matrix models are thus named because they rely on a series of transition matrices. In the forestry sector, the transition matrices express the probability of a forested area leaving its current position within a matrix to join a different position within the matrix, thus acquiring the characteristics (probabilities) of this new category [27]. EFDM is based on the principles outlined in Sallnäs (1990) [28]. EFDM is applied, within this framework, in a manner that reduces data uncertainties traditionally riddling large-scale forestry models related to data input. The philosophy behind EFDM is to involve local partners so that it is possible to use high quality (not necessarily public) data input for the estimation of the current forest state and the transition probabilities from the current state to a future state. The model is configured differently for each national situation in order to reduce the number of assumptions that need to be made when converting data to what is usually a generic configuration for all European countries. The model is implemented in free software precisely to facilitate managing the different locally generated data inputs. In the initial round of computation, EFDM estimates how much wood is “harvestable”. Within this total wood solution, the maximum amount of sawlogs that could be extracted should be specified, and the rest is categorized as pulpwood (the sawn wood potential can then be converted to pulp in GFTM if there is a demand for pulp). Efforts are made to integrate country-specific timber assortment coefficients at this stage, in order to improve estimates for destination commodities. These initial figures are used as a constraint for the production of wood-based commodities and are obtained from the EFDM model at discrete time steps for the duration of the simulation. Based on this constraint, in combination with the exogenous Gross Domestic Product (GDP) trends, the GFTM provides projections of equilibrium prices and quantities produced, consumed, imported, and exported for each commodity and country or group of countries. Input-output coefficients, which in the model transfer input of, for example, logs into sawn wood, express “production efficiency”. They differ between industry branches in various countries, and are therefore instrumental for the model runs. Once equilibrium is achieved in each time step, the GFTM then reports harvest demand from EU forests to the EFDM, divided according to demand for sawlogs and pulp. This division is important as it influences the type of forest to be harvested.

Based on the results of the last iteration in GFTM, EFDM then calculates a provision of woody-biomass. Forest species are often tied to their final product destination. Although the degree of detail in terms of the grouping of species based on their marketable purpose is coarse, it is present nonetheless because a preference can be made in terms of species groups and age groups to be felled in response to the type of demand given by GFTM. Considering species within broad groups also reduces the uncertainties related to the final destination of specific species within different countries. The output from EFDM in this second round of computation will differ with respect to the first round of calculations as the result of adjustments made on the parameters related to forest management.

The modelling flow has so far been described using the EFDM model. However, CBM can be used in the same way in the framework. The CBM is an inventory-based, yield-data driven model that simulates the stand-and landscape-level carbon dynamics of living biomass, dead organic matter and soil, given certain management practices [29]. The model is fully consistent with IPCC methodologies and provides detailed output on all carbon flows in the forest over a given time horizon. Theoretically, data needed to implement CBM and EFDM are very similar, however in practical terms, EFDM makes use of more detailed input data, thus making CBM a viable alternative when detailed data is unavailable.

The Wood Resources Balance Sheet brings all sources and demands of both primary and secondary forest products to a single point, and calculates the excess or deficit. Static in nature, the WRB is simply an accounting tool whose role is to process data from models in the final time step, harmonise the units of data given by all sources, and calculate the initial deficit or excess supply of woody biomass for a specific geographical unit (country or larger region).

### The extended modelling framework

A more complete picture is obtained if the spectrum of tools is extended beyond the core. The energy model POLES [30, 31, 32] processes the proportion of energy from all available sources required in the future, including the proportion generated from the forest sector. In an extended version of the forest-based bioeconomy modelling framework, the energy requirements from the forest sector, as they are forecasted given econometric and climate change information in POLES, are then delivered to the forest sector model GFTM. This total requirement of energy from the forest-based sector may in part be satisfied by secondary sources of wood, such as recycled products. In this case, the total demand for the energy sector must be filtered in way to give a demand to GFTM that is relative to the demand of primary sources (roundwood) to satisfy energy requirements. Secondary sources such as recycled wood must therefore be deducted from the total demand. This data is filtered through the WRB, where the correction coefficients for energy generated from different sources is considered.

In an extended version of the modelling framework, ownership behaviour may be taken into account. This model would influence the amount of wood mobilized based on owner decisions. EVA (Expected Value Asymmetries), is a tool for policy analysis under ownership heterogeneity and may interface between GFTM and EFDM or CBM, in order to simulate the choices made by forest owners [33, 34].

Another important addition to this framework is a land use model. This spatially-explicit model takes into account the competition with other land uses and has the capacity to compute highly spatially-relevant ecosystem services and environmental indicators. The land-use modelling platform LUISA can accommodate sector-specific land requirements, including those of the forestry sector, and resolve the spatial arrangement of land-uses at fine scale, thus taking into consideration competition for land [35, 36, 37]. LUISA attempts to achieve an optimal land-use distribution, based on spatially varying local suitabilities for competing land-uses, while dealing with multiple concurrent EU policies and themes, such as, for instance, transport, urban dynamics, landscape and ecosystem services. EFDM or CBM would provide LUISA with detailed information on the forest structure (age, volume class, species group etc), to compute the appropriate indicators.

Extending the modelling framework is not without disadvantages. The level of uncertainty accumulates, particularly when global-level models that depend on exogenous datasets (that are, in turn, derived from other exogenous datasets) are used. Climate change estimates, population growth estimates and human behaviour have an exceedingly high level of associated uncertainty. Each model has its own mechanism to deal with minimising this inevitable propagation of uncertain data (Lavalle et al. [37]; or a discussion on matrix models in general in Sallnas [28]).

### Hybrid frameworks

On an operational basis, it may not be possible or even desirable to expect data from all components of the framework as described in the previous section. It may be more realistic to expect the core set of tools to interact with one or two other tools. When this occurs, elements of the core system are replaced by more specialized models. For example, the management component of EFDM is replaced by EVA; the contribution of POLES eliminates the need for GFTM to base assumptions about future energy requirements from the forest sector on exogenous proxies; and the contribution of LUISA paints a more accurate picture of environmental impacts of the forest-based sector in the bioeconomy than would direct output from EFDM, et cetera.

When interchanging components (tools) of the modelling system, it is necessary to have a clear idea of the parameters that will be affected with each component addition or removal. The interactions between the tools are described in the following section.

## How the tools interact

Not all parameters of each tool described in Table 1 are exchanged. There are several parameters for the models that can be considered “back-end”, in the sense that they are required to run the models individually, but they are not exchanged with other models or tools in the framework. This section describes the “front-end” parameters, those that are exposed to the other tools in the framework. Table 2 summarises the front-end parameters for each tool.

**Table 2.:**
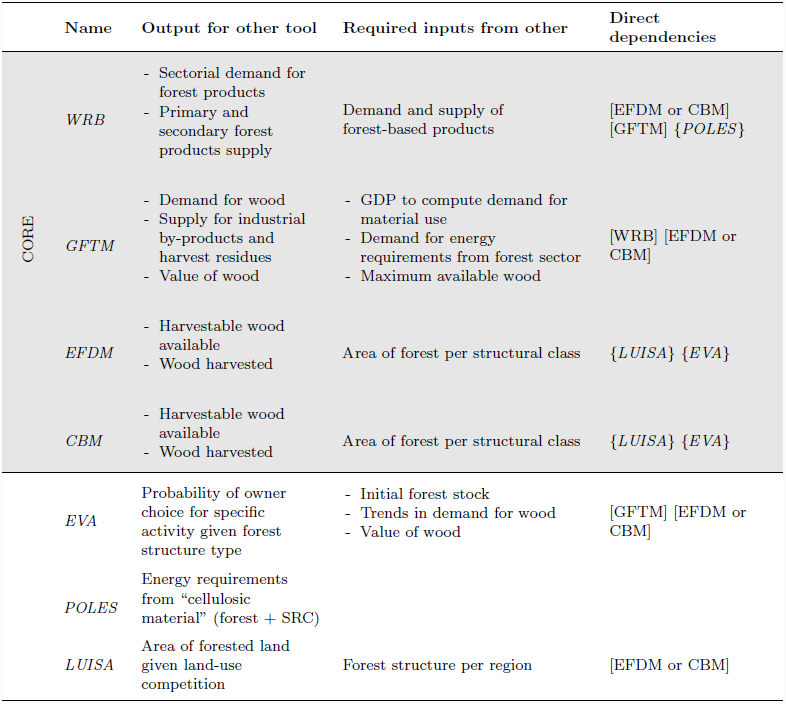
Front-end parameters exchanged within the framework. Optional models are listed in brackets.

To formalize the interactions, variables are assigned to the common data between the tools. Table 3 summarises the interacting variables (front-end), as well as annotations to define common elements.

To avoid errors in computation, integrity checks should exist throughout the system to see if the front-end data is in an appropriate format and that the correct fields are read, otherwise variables can be legitimately and silently read in the system, making it difficult to debug [38]. To ensure a smooth and accurate transition from one model to another, the key intermediate layers of the process are described through a series of concise semantic constraints. The annotation can be made explicit through the use of software language-neutral constraints, described in detail in de Rigo [39]. A first check is that time series contain the same number of rows (::same rows:^2^), thus indicating that the time series is of the same length (T), and must furthermore be of the same division t of finite length θ (::interval:^3^) for different modules within the same simulation must be the same. The time intervals are also contiguous (::contiguous interval:^4^) These rows should be ::numeric:^5^ and contain ::nonnegative:^6^ numbers. It should be verified that the value for final felling per owner group, and forest structure group, is a ::probability::^7^.

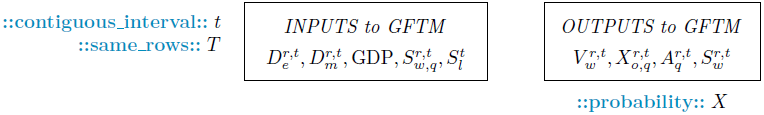

Furthermore, and most importantly, the regional division (*r*) of the data input should be checked. If the regional data is at a finer resolution than the required data, an aggregation needs to be made along the data transfer path. This implies the existence of a look-up table in which all sub-regions are contained within regions (*r* ∈ *R*). For example we can imagine that the data on trends on the initial supply of wood may be given at a small administrative level but for GFTM, a larger region is required. The supply at *r* must therefore be summed to *R*. The system must be aware of what ‘*r*’ are contained within each ‘*R*’ before it can perform the sum. The contrary is also necessary to implement: If the energy model provides a request for energy from the forest sector at national level, this request should be linked to availability at regional level (although trade is permitted, this is important for bookkeeping). Figure 2 shows and example of a subset of data required by GFTM, in a format whereby the integrity check constraints for data input to GFTM needs to be passed before the data is accepted by GFTM.

**Figure 2.**
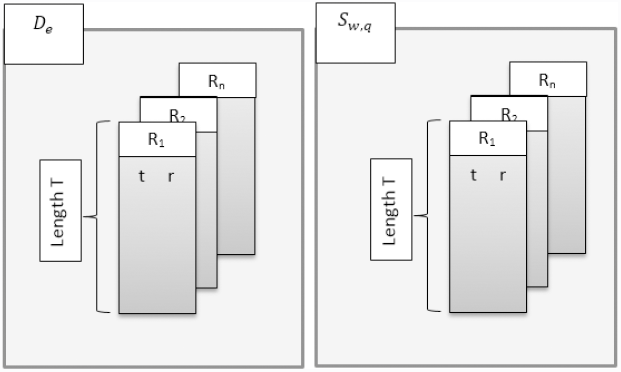
Array-related constraints include number of regions (r) and time steps (t) for the same length of time (T) to process for each input variable.

In Figure 3, the front-end parameters and annotations described n Table 3 can be shown using a sequence diagram where the model interactions are shown in a chronological order, and feedback mechanisms become more clear.

**Figure 3.**
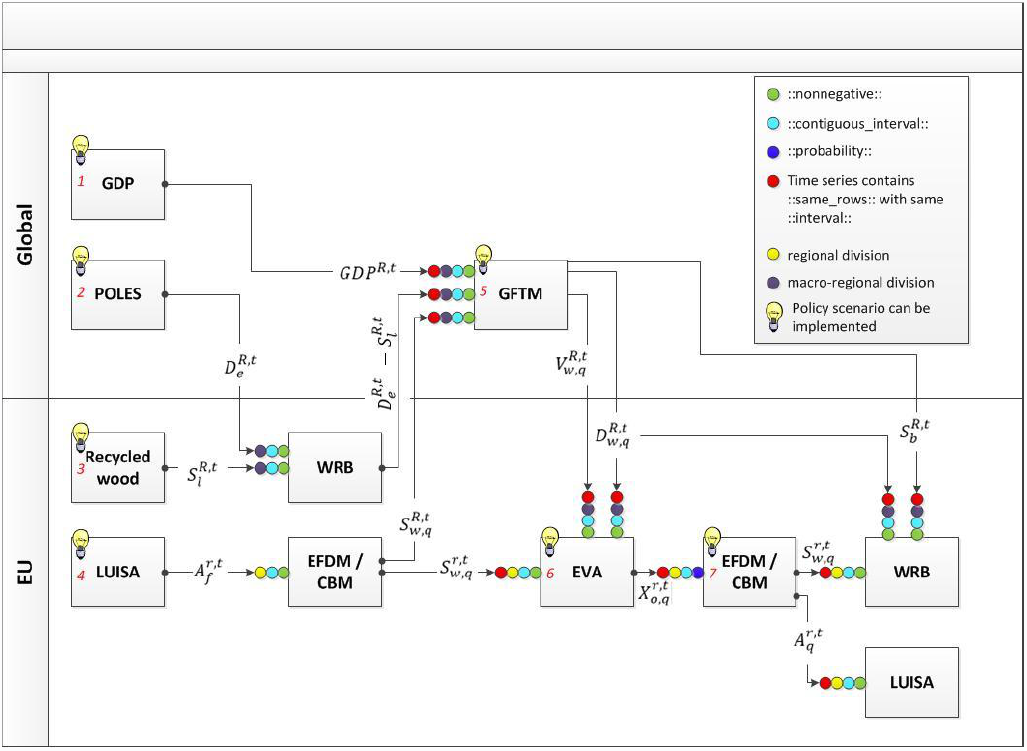
Extended toolset for assessing forest-based sector of the Bioeconomy.

**Table 3.** Interacting variables and annotation between the tools in the Bioeconomy modelling framework

| Variable | Definition | Variable | Definition |
| --- | --- | --- | --- |
| $A$ | Forest area | $S$ | Supply |
| $q$ | Multi-criteria classification of forest structure (e.g. dominant species group, age, volume, stem density, site class etc.) | $w$ | Primary forest products (wood) |
| $f$ | Forest ($q \in f$) | $b$ | Industrial by-products |
| $H$ | Wood harvested | $l$ | Recycled forest products |
| $i$ | Species group ($i \in q$) | $c$ | Secondary forest products (cascading). It is the sum of recycled materials and industrial by-products ($\sum b, l$) |
| $r$ | Simulated region. This can be a $1 \times 1$ km cell or administrative region | $e$ | Energy |
| $R$ | Geographical unit encompassing several regions | $m$ | Material |
| $t$ | Time step class of finite $\theta$ length interval | $V$ | Value |
| $T$ | Final time step of simulation | $X$ | Deterministic probability (0 or 1) of making management decision $o$ for state $q$ |
| $D$ | Demand | $o$ | Owner's decision choice |

The communication between the policy-maker or policy documents and the modeller are not always self-evident. This may partially be the reason why modelling exercises are often criticized for implementing scenarios that are driven by the technological possibility to run the scenario rather than the real questions policy-makers have in mind [40].

The modelling framework described in this paper has been designed in a modular form in order to incorporate the relevant policy with relative ease in different parts of the modelling system. The advantage of the modular system is that of allowing the freedom of assumptions in upstream models to propagate throughout the system, or to replace upstream models whose assumptions may be inconsistent with the scenario configured in a downstream model. The light bulbs shown in Figure 3 indicate where the scenario could act, thus influencing a chain of modelling parameters. To keep track of the parameters that are influenced by the scenario, a denotation that describes the scenario is necessary. The denotation should be linked to a full text description of the scenario, including how parameters are configured to reflect the scenario. In Table 4, the adjusted variable as the result of a policy scenario is denoted by a prime (‘).

**Table 4.**
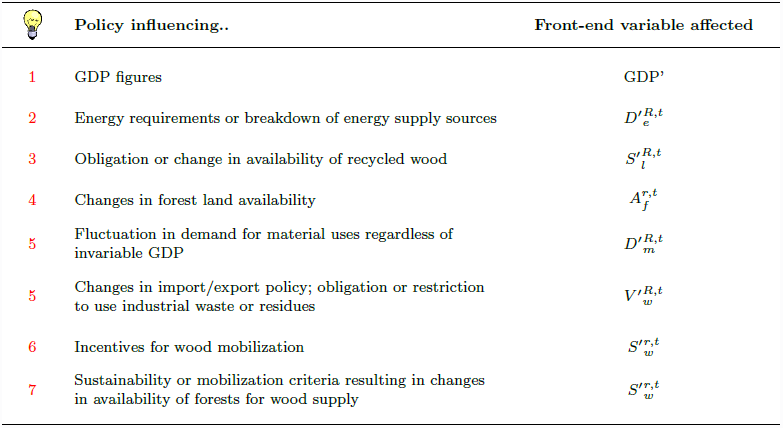
Front-end variables affected by policy scenarios.

|  | Policy influencing.. | Front-end variable affected |
| --- | --- | --- |
| 1 | GDP figures | GDP' |
| 2 | Energy requirements or breakdown of energy supply sources | $D_e'^{R,t}$ |
| 3 | Obligation or change in availability of recycled wood | $S_l'^{R,t}$ |
| 4 | Changes in forest land availability | $A_f^{r,t}$ |
| 5 | Fluctuation in demand for material uses regardless of invariable GDP | $D_m'^{R,t}$ |
| 5 | Changes in import/export policy; obligation or restriction to use industrial waste or residues | $V_w'^{R,t}$ |
| 6 | Incentives for wood mobilization | $S_w'^{r,t}$ |
| 7 | Sustainability or mobilization criteria resulting in changes in availability of forests for wood supply | $S_w'^{r,t}$ |

### Conclusions and way forward

The integration of a partial equilibrium model for the forest sector with a forestry dynamics model and a wood resource balance sheet constitute the core modelling framework for the assessment of the forest-based component of the EU bioeconomy. This framework is expandable to include other sector-specific components, such as the energy model, a land-use model and a model to estimate forest-owner decisions. When integrating the components, several modellers must dialogue. Given that these modellers have different backgrounds, a clear and unambiguous language is required to improve the flow of information. In this paper, we do not discuss the sub-model specific requirements for data, but rather the front-end requirements for model interaction. The data transformation modules (D-TM), developed to link the model components so that they are semantically compliant to the system, are further described in Mubareka et al (in preparation [41]).

The choice to maintain the integrity of the individual models complicates the evaluation of global uncertainty associated to the system. The core modelling components of this system are not yet operational, and software uncertainty has not yet been evaluated as they is still in a prototypical phase, with the exception of the singular operational component CBM [42], however an approach will have to be developed to assess the validity of the system as a whole, probably using real-world data as a benchmark. Opening the system in as much as possible to the public will also allow for a community evaluation of the structure, thus increasing its validity and transparency. While it is true that maintaining the integrity of the individual models complicates the free exchange of the full models or data, the transparency of the system is not necessarily compromised because the interactions between the models through the D-TMs will be made available and will give insight to the system as a whole. Furthermore, the data related to the front-end variables described in Table 3 could be made public. This data, combined with highly transparent documentation on the “black-box” components of the modelling system are added value to the modelling community, and one step closer to encouraging the full openness and transparency of all modelling systems, particularly important for those used in policy support.

## Author Bio

<u>Sarah Mubareka</u> has a Ph.D. in Remote Sensing from the University of Sherbrooke (Canada). She originally joined the Joint Research Center in 2000, and throughout the years has worked on global vegetation mapping, modelling population vulnerability, image processing and land use modelling. Her main area of interest is currently in developing a F/OSS modelling framework for the forestry-based sector of the bioeconomy, to be used to describe the future state of Europe’s forests according to different policy and environmental scenarios.

<u>Ragnar Jonsson</u>’s research focus is on the analysis of forest resources and forest products’ needs at European level, including wood and non-wood products, and the life cycle assessments of these products. He has an M.Sc. in Forestry from the Swedish University of Agricultural Sciences (SLU),as well as an M.Sc. in social sciences from Lund University, and a D.Tech. at Växjö University. As a researcher at SLU, Ragnar produced and analyzed projections of wood products’ demand on European level. His expertise includes the current state of forest management and prospects for increasing the supply of woody biomass within northern Europe. His experience includes econometric analysis of wood-product markets and producing projections of wood products supply and demand at global level for the FAO.

<u>Francesca Rinaldi</u> is a post doctoral researcher whose current research is aimed at studying modeling and analyzing the socio-economic aspects of the forest sector. She holds a degree cum laude in Economics, and a PhD in “Decisions in Insurance and Finance” from the University of Turin. She had previously worked as a research fellow at the University of Turin and at the Manchester Business School. Before joining the JRC she was a researcher of the Financial Economics Research Service of Banque de France.

<u>Giulia Fiorese</u> is a post doctoral researcher whose research focus is on developing an integrated modelling framework capable of assessing greenhouse gas emissions from forestry, agriculture and other land uses and of assessing the impacts of different EU policy scenarios with a specific focus on biomass for energy. Giulia holds a Ph.D. in Ecology from the Universita’ degli Studi di Parma and an M.Sc.in Environmental Engineering from Politecnico di Milano. She has previously worked as a research fellow at Politecnico di Milano and as a research associate at FEEM -Fondazione Eni Enrico Mattei.

<u>Jesús San-Miguel-Ayanz</u> is a senior Researcher at the Institute for Environment and Sustainability of the EC Joint Research Centre in the field of forestry. He holds a PhD in Remote Sensing and GIS from the University of California-Berkeley and a degree in Forest Engineering from the Polytechnic University of Madrid. Prior to jointing the Joint Research Center, Jesús was Associate Professor of Forest Inventory, Forest Mensuration and Remote Sensing and Assistant Professor at the University of Cordoba; and has worked for the European Space Agency and the University of California-Berkeley.

<u>Ola Sallnäs</u> has an M. Sc. in Forestry from SLU (the Swedish University of Agrcultural Sciences), a M. Sc. in Mathematics from Umeå University and a Ph.D. in forestry from SLU. He is at present on leave from his chair in Forest Operations at SLU to work at JRC. His research focuses on forest management planning, forest resource assessment and forest modelling. Lately he has become increasingly interested in national forest policies and their implementation and factual impact on forest management and operations.

<u>Claudia Baranzelli</u> holds an MSc degree in Environmental Engineering with a specialisation in Planning and Environmental Modelling, and a Ph.D in Urban, Regional and Environmental Planning from the Politecnico di Milano. Currently she is scientific officer at the Joint Research Centre where her activities are focused on the development of the Land-Use-based Integrated Sustainability Assessment Modelling Platform. She has experience as researcher in environmental and energy-related modelling, GIS, and planning issues at local and regional level.

<u>Roberto Pilli</u> Roberto Pilli has a PhD in Forest Ecology from the University of Padova. He graduated in Forest and environmental ecology in 1998 and has since worked for the public forest service of the Veneto Region for forest planning and silvicultural activities. He has been providing scientific and technical support on modeling of LULUCF emissions and removals since 2009, when he joined the Joint Research Center and began to work on modeling national forest inventories data, to provide estimates on the forest carbon sink capacity at European and country level.

<u>Carlo Lavalle</u> has over 25 years of experience in applied geophysics and environmental modelling. Since 1990 he is with the Joint Research Centre of the European Commission and has participated in several policies-related Working Groups of the European Union. He is coordinating the development of the Land-Use-based Integrated Sustainability Assessment Modelling Platform.

<u>Alban Kitous</u> is an expert in global energy systems’ modelling and has contributed to numerous international energy and climate policy projects for various institutions. He is now an economic analyst working on modelling for the JRC Unit dedicated to Economics of Climate Change, Energy and Transport (ECCET). Formally educated as an engineer, he also graduated from the Imperial College London MSc in Environmental Technology in 2000.

## Footnotes

1 https://ec.europa.eu/jrc/en/scientific-tool/prospective-outlook-long-term-energy-systems

2 http://mastrave.org/doc/mtv_m/check_is#SAP_same_rows

3 http://mastrave.org/doc/mtv_m/check_is#SAP_interval

4 http://mastrave.org/doc/mtv_m/check_is#SAP_contiguous_interval

5 http://mastrave.org/doc/mtv_m/check_is#SAP_numeric

6 http://mastrave.org/doc/mtv_m/check_is#SAP_nonnegative

7 http://mastrave.org/doc/mtv_m/check_is#SAP_probability

